# Social predation in electric eels

**DOI:** 10.1101/2020.08.10.244129

**Authors:** Douglas A. Bastos, Jansen Zuanon, Lúcia Rapp Py-Daniel, Carlos David de Santana

## Abstract

Social predation, when groups of predators coordinate actions to find and capture prey, is a common tactic among mammals but comparatively rare in fishes. We report the unexpected social predation by electric eels, an otherwise solitary predator in the Amazon rainforest. Observations made in different years and recorded on video show electric eels herding, encircling shoals of small nektonic fishes, and launching joint predatory high-voltage strikes on the prey ball. These findings challenge the hypothesis that electric eels may have a single foraging strategy, and extend our knowledge on social predation to an organism that employs high-voltage discharge for hunting, thereby offering a novel perspective for studies on the evolutionary interplay between predatory and escape tactics.

## Introduction

Social predation occurs when groups of individuals jointly work to find, target and kill larger or more numerous prey ^1^. This foraging tactic is prominently found among mammals, birds, fish and arthropods, and thought to optimize the hunters’ foraging time and energy gain ^2^. Species have long been credited with a conservative and pervasive same set of foraging strategies across populations^1^. However, increasing evidences of populational behavioral diversity challenge this premise^1^. One important behavioral variation among animal populations refers to the use of group foraging as an alternative to single foraging. Group foraging can effectively overcome problems such as detecting, capturing, and controlling prey^2^. Similarly, it can encourage specializations narrowing individual niche breadth and growing resource partitioning among individuals^3^. In turn, the advantages of group foraging may be counterbalanced by competition within the group reducing individual food consumption (e.g. Creel & Creel^4^). That may be the case in situations of resource limitation and intra-populational competition, where groups overtake individuals, but individuals inside groups still compete for limited resource share. Thus, based on model simulations of social structure, Cantor & Farine^5^ proposed a simplified explanation for social foraging to mirror individuals who take decisions to engage in social foraging to attain immediate advantages of foraging with conspecifics to access a common resource. A simple individual-level rule, i.e. taking advantage of catching prey with the help of other individuals, is enough to form temporally stable groups that entirely control the focal food resource. This model takes into account only the current individual experience and does not rely on factors such as actions planning, group structure, past history of membership, or individual relationships. As a consequence, populations can simply display basic prey-use specialization even in the absence of fitness costs or benefits associated with the specific prey.

Despite the possible benefits of group foraging, only few fish species are known to engage in social predation ^1,6^. The electric eels of the Amazon basin have long been considered nocturnal solitary predators, for being capable of employing high-voltage electric organ discharges (EODs) to strike and disable selected prey ^7,8^. One possible explanation to the predominance of lone-hunting by electric eels may be related to the complex behavioral sequence involved in that solitary strategy, which includes prey detection, prey twitch, stunning, and the use of dipole attacks to subdue difficult prey^7^. Thus, this energetically costly but efficient and complex predatory strike would prevent the engagement of other individuals during foraging. Volta’s electric eels (*Electrophorus voltai*) generate up to 860 V during hunting strikes ^9^ on vertebrates and invertebrates ^10^ and are typically observed foraging solo at night, when diurnally-active prey fish are resting in a somewhat lethargic state in shallow waters. Here, we describe the group hunting of Volta’s electric eels, involving over 100 individuals foraging and preying together on shoals of small fishes (Movies S1-S8), which we argue constitutes an unexpected case of social predation ^1^.

## Materials and Methods

During the low-water season, we made field observations near the mouth (maximum 10 meters wide) of a small lake on the banks of the Iriri River (5°34’48.97”S, 54°18’50.95”W). The habitat in which we found the electric eels was structured by sunken logs, with depth ranging from 1.5 to 3-4 meters. The shallow portion of the lake was used as a hunting area, while the deeper portion was used for resting. Limnological parameters were measured in the hunting area in 2014: pH 6.58; electrical conductivity: 20 uS/cm; dissolved oxygen: 5.6 mg/ l; percent of saturation on dissolved oxygen: 15%; water temperature 30.7° C. After euthanizing eight individuals with Eugenol solution, we determined sexes of electric eels by direct gonad inspection (e.g., Waddell & Crampton, 2020). We calculated the approximate maximum distance in which eels stun prey based on video observations. During social predation events, we collected and identified prey and opportunistic predators sharing the lake area. Prey was composed by shoals of small nektonic fishes, mostly characins (*Poptella* spp., *Moenkhausia* spp. and *Tetragonopterus* spp). We recorded one opportunistic predator peacock bass cichlids (*Cichla melaniae*) attacking stunned prey (Movie S7). We estimated prey ball area from still images from the video sequence. We first witnessed the social predation behavior in August of 2012 (Movie S1) and later documented five additional social predation events at the same locality in October 2014, during 72 total hours of continuous observation (Movies S2-S8). We recorded videos with GoPro 3+ and Nikon D5100 cameras. We estimated the number of individual eels involved in the social predation events via direct field observations. To categorize the behavioral states and events, we carried out observations every 30 minutes during the first 24-hour study period to build an ethogram.

## Results

From the 2014 observation, we identified four well-defined behavioral states: 1 – Resting, 2 – Interacting, 3 – Migrating, 4 – Hunting (Figure 1A). This behavioral sequence was witnessed five times consecutively through the entire 72-hour study period. Social predation occurred twice a day. During most of the day (7:30h – 17:00h) and evening (19:30h – 5:00h), male and female adult eels (body length ranging from 1.2 to 1.8 meters) were seen laying almost motionless, close to the mud bottom or among submerged fallen branches and trees at 3 to 4 meters deep. These periods of inactivity were only periodically interrupted by breathing at the surface (Stage 1 - Figure 1B1; 1C1; Movie S2). Around dawn and dusk, eels increased activity, swimming near the water surface and interacting with each other for 20 - 30 minutes at a time (Stage 2 - Figure 1B2; 1C2; Movie S3). On occasion, we watched eels swim together in loose groups spanning ∼20 meters, towards a hunting area that was shallow (<1 m deep) and contained sunken logs that shelter thousands of small fishes (body length: 2-10 cm; Stage 3 - Figure 1B3; Movie S3). During these events, groups of over 100 eels aggregated and starting swimming in circles, herding groups of small fishes into a “prey ball” ^11^ of an area ca. 12 m^2^ (Stage 4 - Figure 1B4; Movie S4). As the herding process progressed, some eels moved into and back from the prey ball (Movie S5) as the rest of the group synchronously pushed it towards a shallower portion of the hunting area. Then, between 2 to 10 individual eels were seen to launch a joint predatory strike, recognizable in our video clips by the conspicuous and synchronized sinusoidal body posture of the striking individuals. Prey hit by the electrical discharges were seen jumping out of the water and returning to the water surface stunned and motionless (Movie S6), being quickly swallowed by the eels or, in some cases, other opportunistic predators (Movie S7). Apparently, the prey ball was attacked each time by different subsets of eels. Each event, including the movement from and to the hunting area, took about two hours from start to end, and involved five to seven joint high-voltage predatory attacks (Movie S8). We propose that this behavior qualifies as a case of social predation.

**Figure 1.**
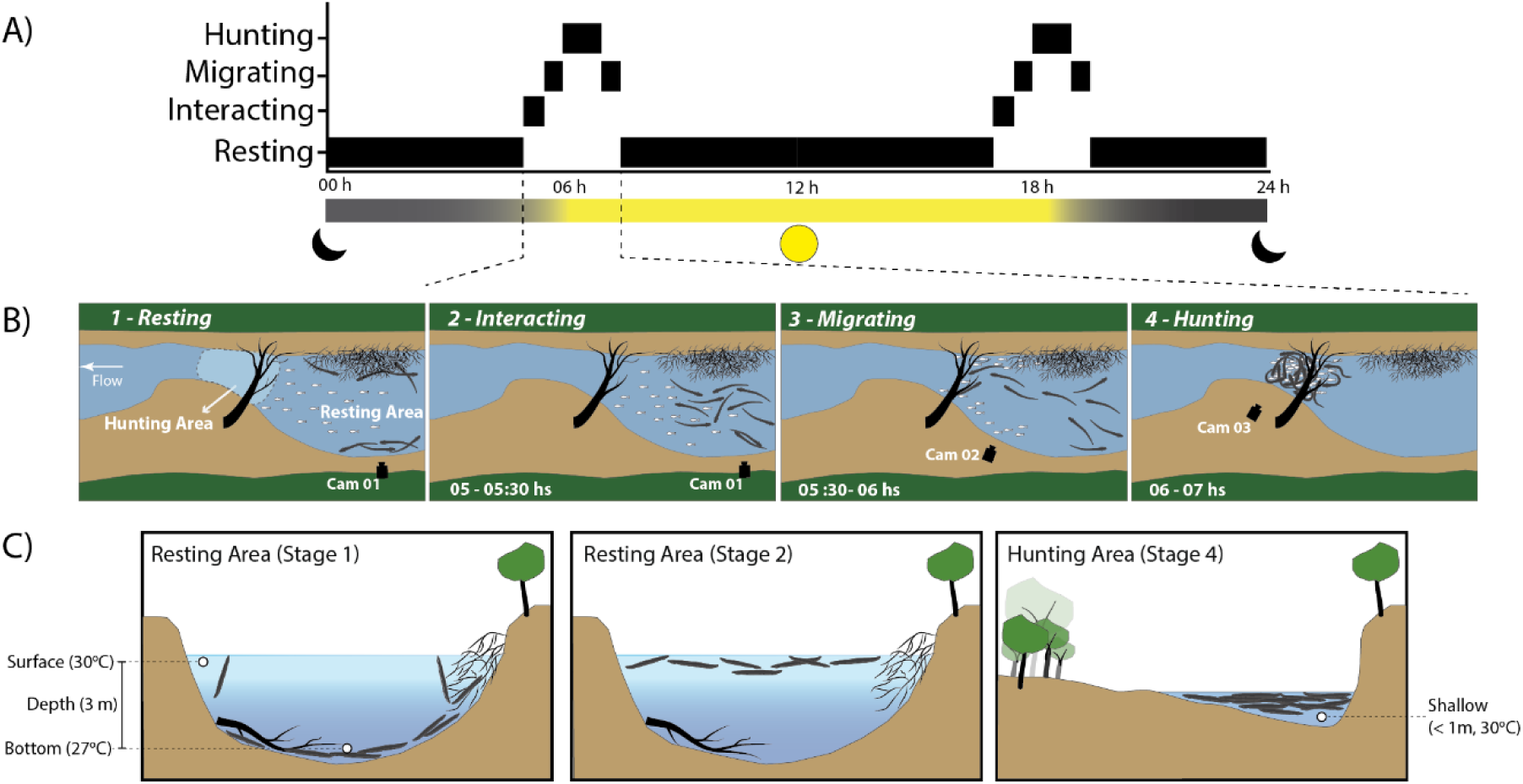
Schematic illustration of stages involved in the social predation as observed in 2014. A) Identified behavioral states throughout 24 hours. B) Aerial perspective during each stage. Stage 1, resting: electric eels were seen laying almost motionless, close to the mud bottom or among submerged fallen branches and trees; Stage 2, interacting: showed increased activity by swimming near the water surface and interacting with each other in the resting area; Stage 3, Migration: group of eels move from the resting area to the hunting area; Stage 4, hunting: groups of over 100 eels aggregate and starting swimming in circles, herding groups of small fishes into a “prey ball”, and posteriorly launching a joint predatory strike. C) Transversal section of the resting (Stages 1 & 2) and hunting (Stage 4) areas; showing the different patterns of spatial occupation by electric eels in the study area and in the water column.

## Discussion

### Social predation in electric eels

Electric eels predominantly prey on single diurnal fish found resting at night in the shallows ^8^, conditions under which ordinary or dipole attacks fired from a very close distance are highly efficient to disable prey ^7^. However, this foraging tactic is probably less efficient if used against shoaling prey during twilight that are aware of the predator’ s presence and can maintain a safe distance. Volta’s eels seem to have overcome these challenges in two ways: by having evolved an increased strength of their highest-voltage EODs ^9^, which may reach and stun prey from relatively large distances (up to ca. 30 cm); and by group foraging ^1,12^ on prey shoals. These two features may assist eels defeating the prey’s anti-predatory responses, i.e. averting from eels and using the confusion effect generated by numerous fish moving amidst a prey ball ^11^. The repeated records of eel groups that performed daily movements between two well-defined places in a same locality across different years are notable, given that individuals were neither confined nor in breeding activity. The low incidence of baseline aggression that we inferred for these large groups of individuals suggests that mutual benefits from social predation may be a driving factor in maintaining these groups of eels^2^.

Thus, we hypothesize that the use of locations with high prey abundance, as well as structural conditions favoring hunting and longtime shelter for multiple eels favor the emergence of social predation by *E. voltai*. Despite been expressed in a multidimensional framework, this social predation could have emerged based on a simple individual-level rule — keep foraging with the same individuals when successful, and that it would suffice for resulting in Voltai’s eels apparent group stability by network self-organization^5^. If this holds true, we expect that social predation events will likely to be registered in other populations living in favorable locations for hunting and resting along *E. voltai*’s distributional range ^9^.

### Electric eels in the context of social predation in a multidimensional framework

The classification of social predation by a single behavioral trait ^1,12,13^commonly fails to fully describe the variety of social predation events found in the animal kingdom^9^. A recently proposed multidimensional framework considers that sociality (1), communication (2), dependence (3), resource sharing (4), and specialization (5) are five dimensions of social predation ^1^. Based on Lang & Farine’s^1^ subclass scoring framework for each of the five dimensions, we recognized the presence or absence of these key features and propose that Volta’s electric eels can engage in social predation events based on our observations that individual eels: (1) social — repeatedly forage and feed together over time; (2) signaling — apparently communicate with each other by active body posturing; and (3) high dependence — are dependent on collective actions when hunting together. However, we note that (4) competition — when hunting in groups, individual eels apparently share prey randomly; and (5) no specialization — do not have well-defined individual roles. Taken together our results strongly suggest that electric eels can be placed in the social predation behavioral landscape ^1^.

### Caveats

Our data have some limitations that impair strictly considering the proximate causes of the observed social predation strategy by *E. voltai*: 1) We don’t have EODs data during eel’s group foraging, which precludes analyzing the role played by low-voltage EODs during intra-specific communication, as well as the use of high-voltage EODs repertory during the social predation events. For instance, we cannot ascertain if electric eels use low-voltage EODs to recruit individuals, nor if they use high-voltage EODs to detect fast moving prey ^4^ and/or to drive prey shoals in the hunting area. More importantly, EOD recordings during social predation events would allow to verify if only a small subset of the eels’ group produce, altruistically, high-voltage strikes benefitting a larger number of individuals from the consumption of the stunned prey.; 2) The absence of genetic data regarding the eels engaged in social predation events^5^. The lack of fine-scale genetic data undermined our capacity to understand possible kin relations and maybe hierarchical structures within the group. Likewise, a broader genetic comparison across populations of *E. voltai* would allow us to infer whether electric eels foraging networks are resilient to stochastic events (e.g. Cantor & Farine^5^); 3) The lack of behavioral data for a quantitative comparison of the foraging success between group and solitary hunting, which would allow us to access whether social predation results in foraging time and energy gains over solitary hunting.

### Perspectives

Despite the limitations aforementioned, our findings advance the knowledge on social predation by extending it to a large vertebrate that employs high-voltage discharges for hunting, showing that those animals have a broader hunting repertory than previously known, challenging the hypothesis that many species have a single foraging strategy^1,5^. In addition to trying to locate additional populations of eels involved on group foraging, our future field and lab-based studies will investigate social predation in electric eels focusing on the link between population, social structures, genomics, and electrogenesis. In short, this case offers a unique perspective for future studies on the evolutionary interplay between predatory and escape tactics among vertebrates.

## Acknowledgments

This research was funded by a National Geographic grant (#9519-14) and Global Genome Initiative (#2017-149). We thank T. de Souza from the Instituto Chico Mendes de Conservação da Biodiversidade for logistical support. DAB and JZ were founded by a Brazilian National Council for Scientific and Technological Development (CNPq) grants (# 140145/2016-8 and #313183/2014-7, respectively). This paper was greatly benefited by the comments of M. Cantor, J. Benda and J. Podos. Thanks P. Bartsch and J. Ziermann for fruitful discussion. CDS was funded by the São Paulo Research Foundation/ Smithsonian Institution grant (#2016/19075-9).

## Supplementary materials for this manuscript include the following

Movies S1 to S8

**Movie S1**: First record of social predation in *Electrophorus voltai* during the low-water season in the mouth of a lake in the Iriri River drainage in 2012. 1). Group of electric eels (adult males and females; body length ranging from 1.2 to 1.8 meters) swimming in the hunting area; 2) A subset of electric eels (ca. 30 individuals) striking and disabling shoals of small fishes.

**Movie S2: Stage 1 – Resting**: Electric eels surfacing to breathe (red arrows). 1) General view of resting area; 2) Close-up of electric eels gulping air, peacock bass cichlids (*Cichla melaniae*) can be seen swimming among eels. Camera 1 (Nikon D5100), real time (Figure 1B1).

**Movie S3: Stage 2 and 3 - Interactions and Migration**: Electric eels swimming near the water surface. 1) Eels intraspecific interaction (duration 20 - 30 minutes); 2) Eels swimming together as a loose group migrating through a stretch of ∼20 meters towards a shallow (<1 m deep) hunting area. 3) Eels migrating to the resting area after coordinated hunting. Camera 1 (Nikon D5100) accelerated x6, Camera 2 (Nikon D5100) accelerated x3 and Camera 3 (Nikon D5100) accelerated x4.5 (Figure 1B).

**Movie S4: Stage 4 - Prey ball**: A prey ball is composed by shoals of small nektonic fishes, mostly characins (*Poptella* spp., *Moenkhausia* spp. and *Tetragonopterus* spp.). 1) Video frames indicating presence (red arrow)/absence of the prey ball (dark blotch) in the hunting area; 2) The shoals of fishes moving, inside the circle, during prey ball formation; 3) The same previous video with the prey ball highlighted (green blotch inside the circle). The effects were applied: black and white, high contrast and Lumeri color (Changing the dark blotch to the green blotch), using Adobe Premiere Pro 2020.Camera 3 (GoPro 3+) accelerated x3 (Figure 1B4).

**Movie S5: Stage 4 – Electric eels herding a prey ball**: Group of electric eels conducting the prey ball to the shallow area. Prey ball estimated from video clips to occupy an area ca. 12 m^2^. The coordinated driving movements are represented by three eels (tracked frame by frame using Adobe After Effects 2020). The green line represents the forward movement, circling or entering the center of the prey ball. The red line represents the reverse movement, usually leaving the center of the prey ball. Camera 3 (GoPro 3+) accelerated x7 (Figure 1B4).

**Movie S6: Stage 4 – Electric eels attacking the prey ball**: The attacks usually start at the margin and extend until the middle of the channel. 1) High-voltage strikes showing preys jumping and falling stunned on the water surface; 2) High-voltage strikes inside the prey ball can be identified by the sinusoidal body posture (highlighted). Camera 3 (GoPro 3+) real time (Figure 1B4).

**Movie S7: Opportunistic predators (peacock bass cichlid**): Opportunistic behavior of peacock bass cichlids (*Cichla melaniae*) during coordinated hunting in electric eels. 1) Peeking at the prey ball; 2) Attacking stunned preys; 3) Peacock bass jumping when stroked by high-voltage discharges. Camera 3 (GoPro 3+) real time (Figure 1B4).

**Movie S8: Stage 4 – Full coordinated hunting event:** Accelerate video from start to finish of coordinated hunting behavior (Stage 4): electric eels arriving at the hunting area (18h). Then, the eels begin herding and pushing the prey ball to the shallows. Different subgroups (ca. 5 to 50 individuals) make a series of five attacks between 18:17 and 18:35 h. After the attacks the eels migrate back to the rest area at dusk (18:40 h). Group hunting occurred twice a day. Camera 3 (GoPro 3+) accelerated x41.8 (Figure 1B4).

